# Individual Learning Phenotypes Drive Collective Cognition

**DOI:** 10.1101/761676

**Authors:** Chelsea N. Cook, Natalie J. Lemanski, Thiago Mosqueiro, Cahit Ozturk, Jürgen Gadau, Noa Pinter-Wollman, Brian H. Smith

**Author notes:** Co-Senior Authors (Noa Pinter-Wollman, Brian H. Smith). **Author Contributions**: CNC, TM, JG, NPW, BHS conceived of and helped design the study. CNC, NJL, and TM analyzed the data. CNC and CO created the genetically selected lines and CO maintained them. CNC carried out data collection and wrote the first draft of the manuscript. CNC, TM, NJL, NPW, BHS discussed results. All authors commented on the manuscript.

## Abstract

Variation in cognition can influence how individuals respond to and communicate about their environment, which may scale to shape how a collective solves a cognitive task. However, few empirical examples of variation in collective cognition emerges from variation in individual cognition exist. Here, we show that interactions among individuals that differ in the performance of a cognitive task drives collective foraging behavior in honey bee colonies by utilizing a naturally variable and heritable learning behavior called latent inhibition (LI). We artificially selected two distinct phenotypes: high LI bees that are better at ignoring previously unrewarding familiar stimuli, and low LI bees that can learn previously unrewarding and novel stimuli equally well. We then provided colonies composed of these distinct phenotypes with a choice between a familiar feeder or a novel feeder. Colonies of high LI individuals preferred to visit familiar food locations, while low LI colonies visited novel and familiar food locations equally. However, in colonies of mixed learning phenotypes, the low LI bees showed a preference to visiting familiar feeders, which contrasts with their behavior when in a uniform low LI group. We show that the shift in feeder preference of low LI bees is driven by foragers of the high LI phenotype dancing more intensely and attracting more followers. Our results reveal that cognitive abilities of individuals and their interactions drive emergent collective outcomes.

**Significance Statement:** Variation in individual cognition affects how animals perceive their environment and which information they share with others. Here we provide empirical evidence that how individual honey bees learn contributes to collective cognition of a colony. By creating colonies of distinct learning phenotypes, we evaluated how bees make foraging choices in the field. Colonies containing individuals that learn to ignore unimportant information preferred familiar food locations, however colonies of individuals that are unable to ignore familiar information visit novel and familiar feeders equally. A 50/50 mix of these phenotypes prefer familiar food locations, because individuals who learn the familiar location recruit nestmates by dancing more intensely. Our results reveal that variation in individual cognition scales non-linearly to shape collective outcomes.

## INTRODUCTION

Collective behavior allows animals to undertake tasks that they could not accomplish alone. Individuals utilize local information to adjust to ecological changes as a collective. Local information is implicitly or explicitly communicated among group members to form a collective response (1–3). Individuals within a group vary in their cognitive abilities. Cognition at the individual level occurs when an organism perceives, integrates, and utilizes acquired information. Collective cognition is a form of collective behavior that emerges from the interactions among individuals working together to solve a cognitive task that could not be accomplished as effectively at the individual level (1, 4). Many of the basic rules that explain collective behavior and cognition come from theoretical modes, which emphasize the importance of variation in perception and cognition among individuals within a social group (5). For example, leaders can emerge in computer simulations to guide uninformed group members to a resource. However, both informed and uninformed individuals are needed to effectively move in the correct direction (6). Although individual variation in responsiveness and cognitive ability is recognized as critical for the emergence of collective cognition, empirical work on the mechanisms by which variation in individual cognition and the interaction between these different behavioral types scales to the collective are rare.

One way in which animals differ from one another in their cognitive abilities is the way in which they perceive information (7). This perception may be driven by several cognitive properties, including the ability to learn relevant information. This ability has important ecological and evolutionary consequences(8). For example, learning is the foundation of the evolution of aposematic coloration (9). Humans that are able to quickly learn important information report increased productivity compared with individuals that cannot focus on pertinent information (10–12). Naturally, collective groups of organisms will consist of individuals that vary in how they learn information. Here we ask how individual variation in learning shapes the way in which individuals learn and share ecological information with group members to shape collective outcomes.

While foraging, honey bees (*Apis meillifera*) must learn various aspects about the location of food sources, such as landmarks, odors, and direction (13–15). Honey bee foragers then return to the colony to communicate this spatial information to colony members at the nest via their recruitment dances(13). In the lab, honey bees exhibit variation in their ability to learn to ignore unimportant information, such as unrewarding odors, known as latent inhibition (16, 17). LI has been studied in vertebrates (18–22) and is correlated with attention disorders in humans (10). LI is heritable in honey bees (23). Foraging honey bees vary in their expression of LI; scouts tend to exhibit high LI and ignore familiar odors, while recruits tend to exhibit low LI and learn familiar and novel odors equally well (24). Despite our knowledge of variation among individuals in latent inhibition (23, 25), and its effects on predator avoidance (18, 19, 26), it is unknown whether or how this variation affects ecologically relevant decisions in social systems.

We provide empirical evidence that the interaction of individuals that vary in their cognitive abilities drives collective cognition. Using the genetic heritability of LI, we first tested reproductive queen and drone honey bees to characterize their LI, then we selected two distinct phenotypes from the reproductive individuals: high LI and low LI. We then created genetic learning lines from singly inseminated queens by like performing drones to produce two distinct lines of workers that exhibit similar LI to their parents. First, we verify that the social environment of adult honey bees from selected lines does not affect their LI phenotypes as foragers. We then created 24 colonies composed of single cohorts of only low, only high, 50/50 mixed high and low LI workers, as well as age-matched non-selected control bees. To compare collective foraging behavior across these selected colonies, we placed them in semi-natural foraging conditions, then evaluated the number of forager visits, first visits, and re-visits to the familiar or novel food locations. To explore the mechanisms underlying how individual variation in LI affects collective foraging, we quantified the round recruitment dance in 6 mixed colonies while the colonies visited novel and familiar feeders. These experiments allowed us to simultaneously quantify how collectives vary in performing cognitive tasks as a result of the composition of the individuals of that collective, as well as how cognitively distinct individuals interact to shape collective outcomes.

## RESULTS

To ensure workers in different social environments exhibited the predicted heritable LI phenotype, we evaluated the LI score of foragers after 21 days in either their natal colony or a control colony. We marked 1000 individuals from each selected line (high or low LI) on the day of emergence. We then placed 500 individuals back into their natal colony and 500 individuals into an established control colony of equal size with an open mated queen, i.e. workers with a variety of learning phenotypes. We monitored the colonies until marked bees began to make foraging flights (∼21 days). We then collected marked foragers as they returned to the colony and brought them into the laboratory to evaluate their LI. We avoided pollen foragers as they tend to exhibit different learning behavior compared to nectar foragers(27). We found that foragers retained the expected LI based on the LI of their parents, regardless of whether they were housed with same or with variable learning phenotypes. Foragers from the high and low lines differed in expression of LI as expected (GLM: χ^2^ = 4.84, df=1, p=0.027, Figure 1). We did not detect an effect of the identity of the colony in which the bees were housed on LI phenotype (χ^2^ = 3.28, df=2, p=0.193, Figure 1).

**Figure 1:**
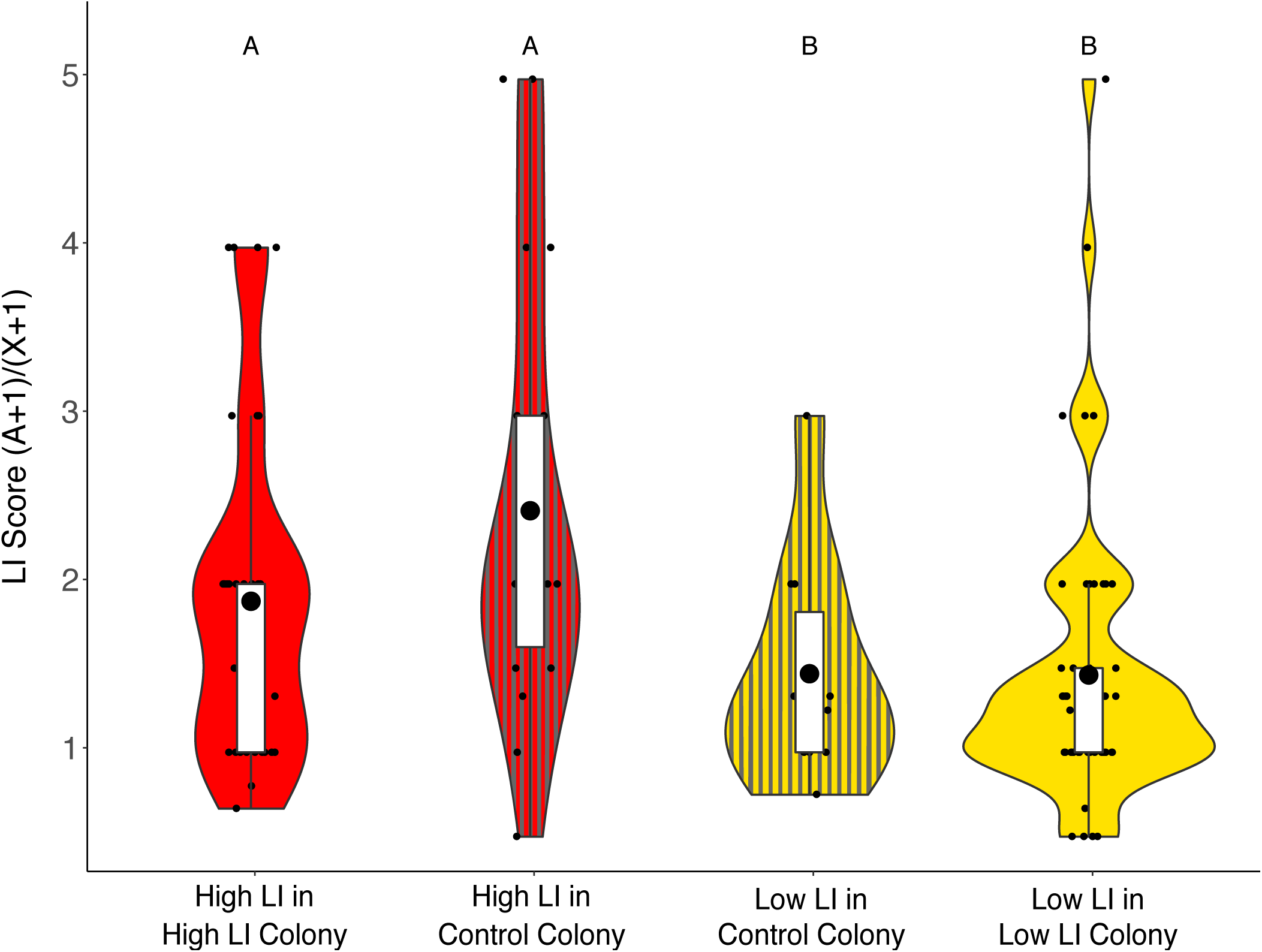
Social environment does not alter expression of genetically selected latent inhibition. LI scores of individuals from high LI lines that spent their adult life either in high LI only colonies (red, n=36) or in a control colony with a variety of LI phenotypes from an open mated queen (red with gray vertical lines, n=18); individuals from low LI lines that spent their adult life either in low LI only colonies (yellow, n=52) or in control colonies (yellow with gray vertical lines, n=10). In this and subsequent figures, the large black dot is the mean, the white box is the interquartile range (IQR), whiskers extend to 1.5*IQR, and the small points beyond the whiskers are outliers. Shaded areas show the distribution of the data. Here, and in all following figures, yellow are low LI colonies and individuals, gray are control colonies and individuals, and red are high LI colonies and individuals. Attention is critical for many individual behaviors, including finding the correct mate or prey.

To determine how the learning phenotypes influenced colony-level foraging behavior, we placed small single-cohort (same age bees) colonies into a flight cage and monitored foraging activity. We evaluated 4 colony types each week: one control colony consisting of approximately 1300 age-matched bees from open mated queens; one colony consisting of 650 workers from high LI queens plus 650 age-matched control bees; one colony consisting of 650 workers from low LI queens plus 650 aged-matched control bees; and one 50/50 mixed colony with 325 workers from each LI line plus 650 aged-matched control bees. In the last 3 types, the supplemented 650 age-matched bees from open mated queens were used to ensure a small but functioning colony as we did not have enough workers from the single-drone-inseminated queens and colonies of just 650 individuals would be too weak to forage. Honey bee division of labor is largely influenced by worker age, so we used age-matched bees to remove any influence that age may have on foraging propensity. On day 1, we trained bees to a feeder inside the tent containing 1M sucrose and an odor, which became the ‘familiar’ feeder. During the subsequent 3 days, in addition to the familiar feeder, we introduced a single novel feeder each day with a different odor and color, but with the same sugar concentration as the familiar feeder (Figure 2A). To evaluate the collective ability of the colony to find a new feeder, we recorded the number of visits to each feeder by bees from each selected line according to the color of paint on the bees’ thorax. We further marked bees with a feeder-specific color on their abdomen when they visited the feeder for the first time to determine if bees revisited that feeder. We repeated this for 6 weeks on 6 colonies for each group type.

**Figure 2:**
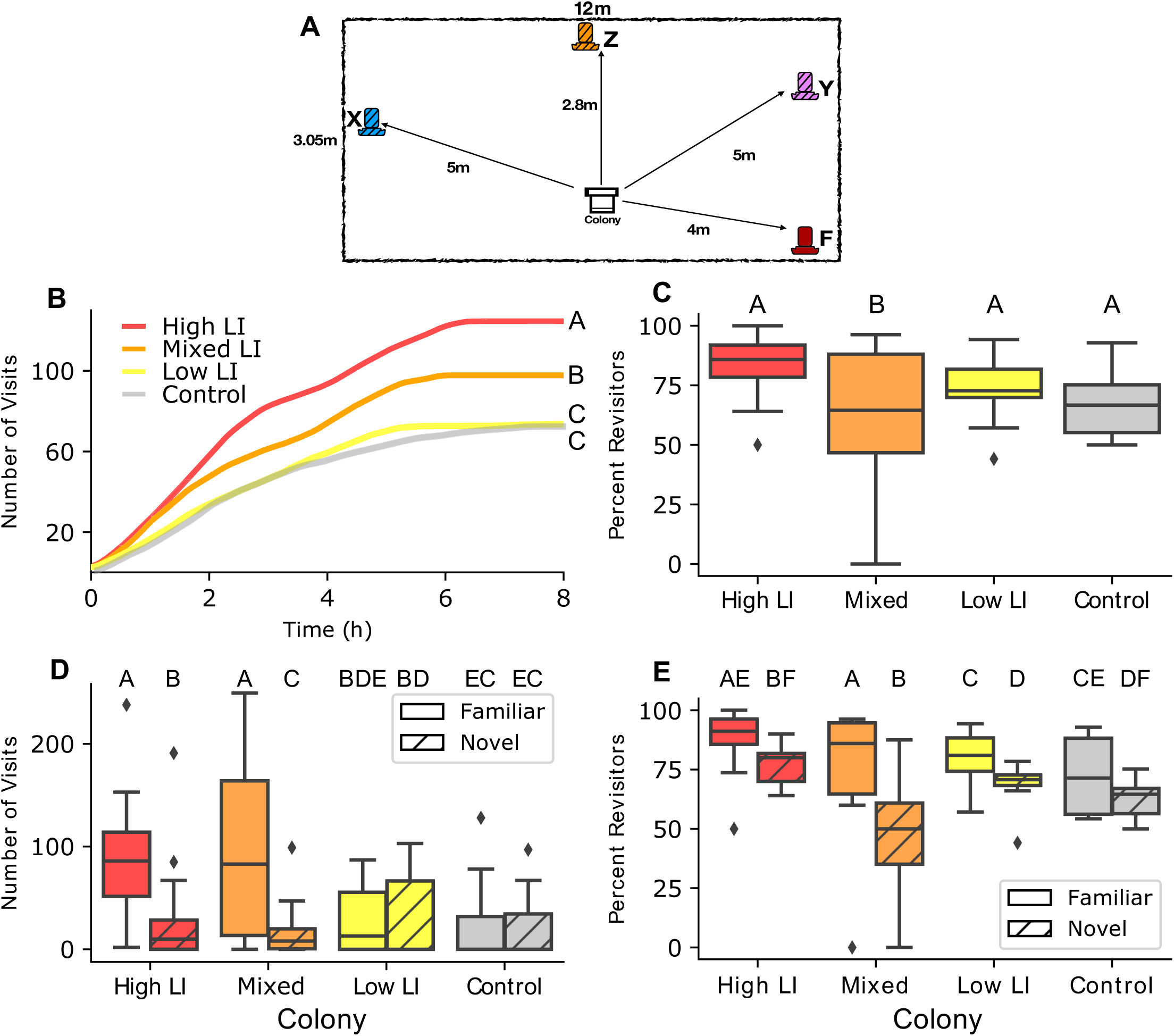
Colonies constructed from different genetic lines selected for high or low latent inhibition exhibited differences in collective foraging behavior. (A) The experimental set up illustrating the location of feeders in relation to the location of the colony (center, white) within the experimental arena (large rectangle). The familiar feeder (red) was provided on day 1 and on all subsequent days. Novel feeder X (blue) was presented on day 2, novel feeder Y (purple) on day 3, and novel feeder Z (orange) on day 4. See Supplementary table 2 for associated odors. Visits to all novel feeders were combined for statistical analysis. (B) Cumulative number of visits of bees to all feeders over time by colony type. Different letters to the right of the lines indicate statistically significant differences according to a post hoc Tukey test. For further illustration of visitation by each colony on each day, see Supplementary Figure 1. (C) Percent of re-visits out of the total number of visits to all feeders by colony type. Here and in all following panels, different letters above boxes indicate statistically significant differences according to a post hoc Tukey test. (D) Number of all visits to the familiar feeder (solid boxes) and a novel feeder (hatched boxes) for each type of colony, when both novel and familiar feeders were presented simultaneously (days 2-4). (E) Percent of re-visits out of the total number of visits to either the familiar or the novel feeder by type of colony when both novel and familiar feeders were presented simultaneously (days 2-4). In C, D and E, horizontal lines are the median, the boxes are the interquartile range (IQR), whiskers extend to 1.5*IQR, and the small points beyond the whiskers are outliers. N=24 colonies, 6 colonies per group type, 6172 total visits.

Colony composition strongly influenced overall number of visits to the food locations (N = 6 colonies in each line, 24 total, 6172 total visits; GLM: χ^2^ = 1270, df = 3, p < 0.0001, Figure 2B). High LI colonies had significantly more visits to all food locations compared to low LI colonies (Tukey post hoc: Z=25.5, p <0.0001, Figure 2A), mixed colonies (Z=5.18, p<0.0001), and controls (Z=26.6, p<0.0001). Mixed LI colonies also had significantly more visits compared to low (Z=20.7, p < 0.0001) and controls (Z=21.8, p<0.0001). Low LI and control colonies had the fewest total visits and were not significantly different from each other (Z=-1.38, p=0.50).

Foraging in the high, low, and control colonies was largely performed by bees revisiting the feeders. (GLM, χ^2^ = 22.32, df =3, p<0.0001, Figure 2C). However, the mixed LI colonies had a significantly lower proportion of revisiting foragers compared to the low (Tukey post hoc: Z=-4.2, p=0.0002), high (Z=-3.1, p=0.01), and control colonies (Z=-3.33, p=0.004). We did not detect significant differences among the other colony types (See Supplementary Table 3).

A colony’s LI phenotype composition determined its preference between the novel and familiar feeders (GLM: Feeder*Colony χ^2^ =473.64, df=3, p<0.0001; Figure 2D). High and mixed colonies preferred the familiar feeder over the novel one (Tukey Posthoc: High Familiar:Novel: Z=20.2, p<0.0001; Mixed Familiar:Novel: Z=25.6, p<0.0001). Low LI and control colonies did not show a strong preference for either feeder, visiting them equally (Low Familiar:Novel: Z=-1.24, p=0.92; Control Familiar:Novel: Z=2.03, p=0.46). For full pairwise comparisons, see Supplementary Table 4).

The number of re-visits to the novel and familiar feeders was different across colony compositions (Figure 2E: Colony*Feeder χ^2^ =53.67, p<0.0001). All colonies had a higher proportion of re-visits to the familiar feeder compared to the novel feeder. However, the mixed LI colonies had a much lower proportion of re-visitation to the novel feeders than the other colony types (Supplementary Table 5). Thus, new foragers in the mixed colonies that visited the novel feeder were less likely to return to it compared to foragers who visited the novel feeders in other colonies.

To determine why the mixed colonies showed a preference for the familiar feeder (Figure 2D), we examined how individual lines visited each feeder (Figure 3). In 2017, we tested mixed colonies placed in a flight cage. In 2018, we reselected lines and then placed mixed colonies into two-frame observation hives to evaluate recruitment dances along with visitation to the feeders in the flight cages. We found that there was a significant year effect (Supplementary Table 6), likely due to reselection and different environmental conditions. We therefore statistically analyzed each year separately to focus on the within-year variation between the selected lines.

**Figure 3:**
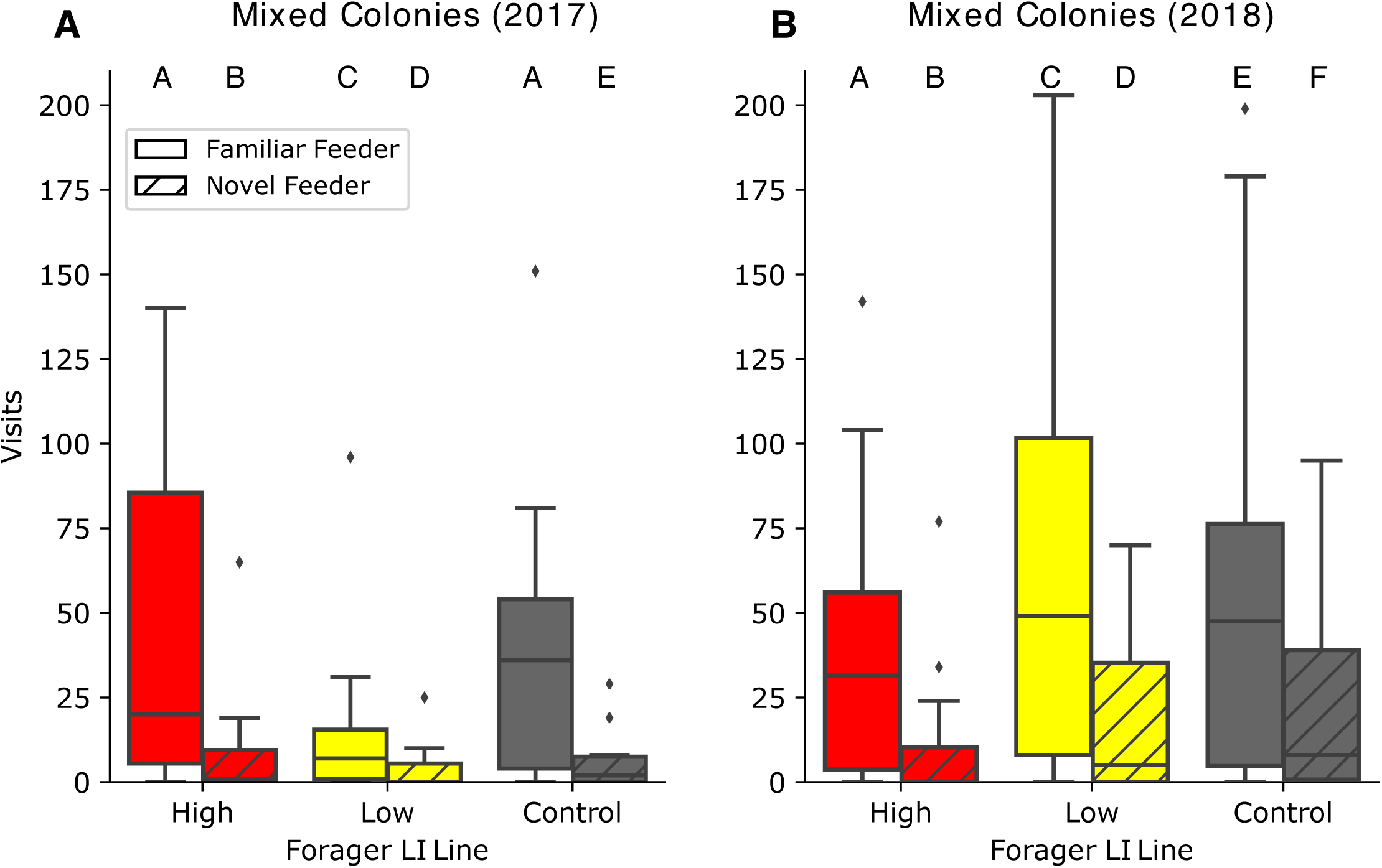
Visits of individuals from different genetically selected lines when in a mixed colony. Daily visits to the familiar (solid) and novel (hatched) feeders by individual bees in mixed colonies from low LI parents (yellow), high LI parents (red) or open mated queens (grey) in (A) 2017, N=6 mixed colonies, 2347 overall visits and (B) in mixed colonies from lines that were re-selected in 2018, N=6 colonies, 6272 overall visits. The horizontal line in the box is the median, the box is 25-75% of the data, whiskers represent 95% of the data, and diamonds show outliers beyond 95%. Different letters above boxes indicate statistically significant differences according to a post hoc Tukey test.

Low LI and control individuals shift their preference to the familiar feeder when mixed with high LI bees. In 2017, we found a significant interaction between the selected line and which feeder foragers visited (GLM: χ^2^ =7.79, df=2, p=0.02; Figure 3A). Although low LI and control colonies did not show a preference to a novel or familiar feeder when they had a uniform colony composition (Figure 2E), when mixed with high LI individuals, low LI and control individuals exhibited a preference to the familiar feeder (GLM: Low Familiar:Novel: Z=13.28, p<0.0001; Control Familiar:Novel: Z=18.32, p<0.0001; Figure 3A). High LI individuals showed a preference to familiar feeders (GLM: Familiar:Novel: Z=22.03, p<0.0001) just as colonies comprised of only high LI individuals did (Figure 2E). We found a significant interaction between selected line and feeder in 2018 (GLM: χ^2^ =85.27, df=2, p<0.0001; Figure 3B), with low LI and control individuals showing preference to the familiar feeder over the novel feeder (GLM: Low Familiar:Novel: Z=25.05, p<0.0001; Control Familiar:Novel: Z=13.90, p<0.0001; Figure 3B) similar to high LI individuals preferring the familiar feeders (Familiar:Novel: Z=18.48, p<0.0001). For full contrasts from 2017 see Supplementary Table 7, for 2018 see Supplementary Table 8.

To uncover the behavioral mechanisms that underlie the switching of low LI individuals from having no feeder preference when in a uniform colony composition to preferring the familiar feeder when in a mixed colony, we examined the round dance, the modified waggle dance used at short distances(28), of individuals from each selected line in mixed colonies as they returned from foraging. Using observation hives with glass walls, we video recorded bees performing the round dance to recruit other individuals in the colony to forage. To determine which selected line recruited to each feeder, we noted the selected line of the dancer (high or low LI) according to the paint marks on the individuals’ thorax and whether the dancer had visited a feeder according to the paint marks on abdomens. We did not record dancers without abdominal marks as they were likely collecting from and recruiting to unmonitored water sources. To determine who the information about a feeder was communicated to, we counted the number of followers of each dancer and the selected line of the followers. To quantify the dance intensity, we recorded the duration of the dance, and the number of turns the dancer made during the first 20 seconds of the dance.

Individuals from the lines differed in their likelihood to perform a round dance (Chi-square test: χ^2^=26.61, df=2, p<0.0001; Figure 4B). Low LI individuals were significantly more likely to perform a dance compared to high Li individuals (pairwise chi-square test: p=0.0001) and controls (pairwise chi-squared test: p=0.004). High LI individuals were just as likely to perform a dance as controls (p=0.36). Individuals differed in their likelihood to follow a dance based on their selected line (Chi-square test: χ^2^=28.26, df=2, p<0.0001; Figure 4B). Low LI individuals were significantly more likely to follow a dance compared to high LI bees (pairwise chi-square test: p<0.0001) and controls (pairwise chi-square test: p<0.003). High LI and control individuals were equally likely to follow a dance (pairwise chi-square test: p=0.240).

**Figure 4:**
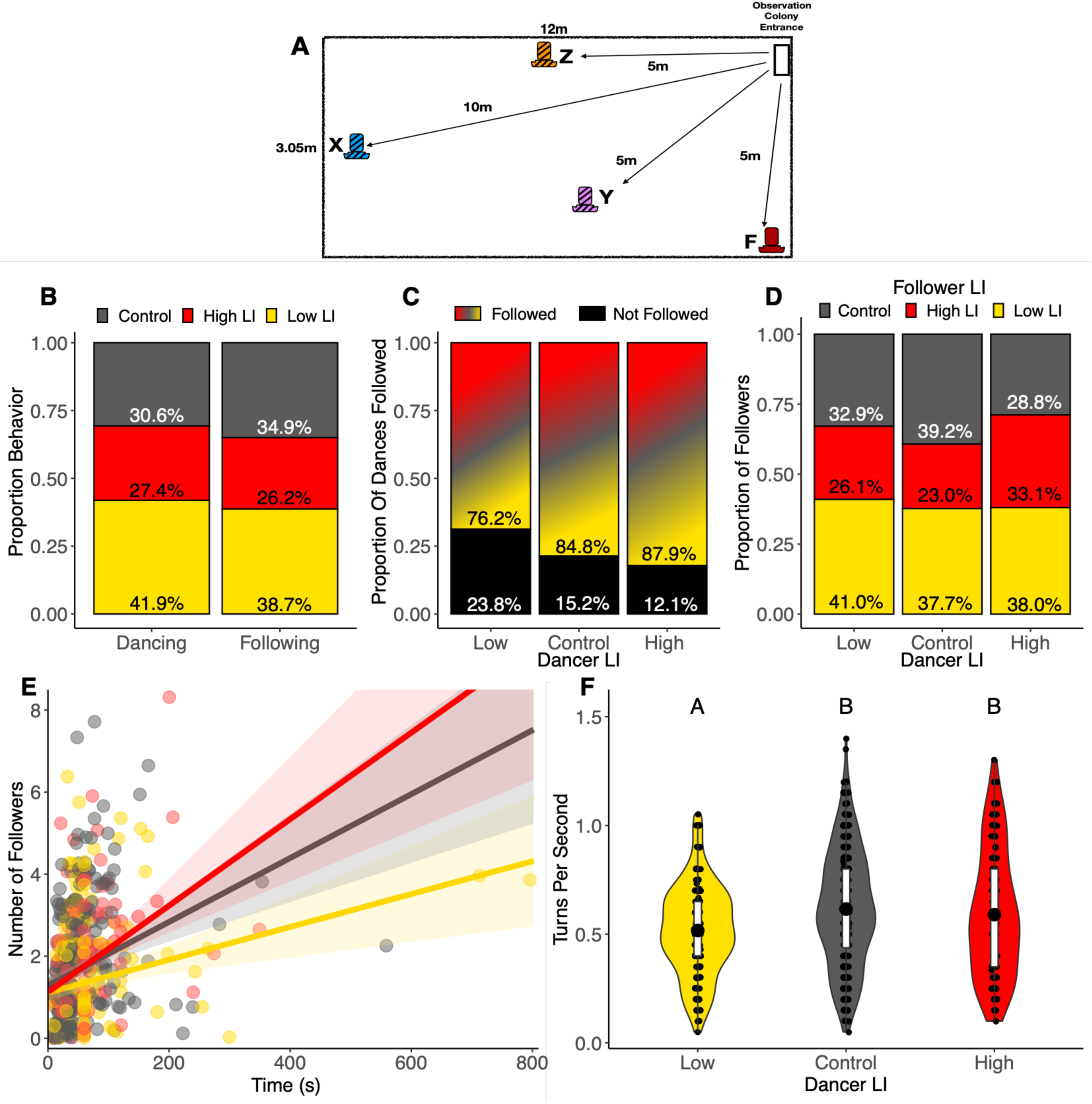
Recruitment dances facilitate integration of information from different genetically selected lines. (A) The experimental set up illustrating the location of feeders in relation to the location of the colony entrance (top right, white) within the experimental arena (large rectangle). The familiar feeder (red) was provided on day 1 and on all subsequent days. Novel feeder X (blue) was presented on day 2, novel feeder Y (purple) on day 3, and novel feeder Z (orange) on day 4. See Supplementary table 2 for associated odors. Visitation to novel feeders were combined for statistical analysis. (B) Proportion of dances (N=667) or follows (N=1201) across 6 colonies performed by bees from each line, relative to their abundance in the mixed colony (350 high, 350 low, 700 control). We accounted for the difference in abundance of each selected line by dividing the number of observed control dancers by 2 before calculating these proportions. (C) Proportion of dances performed per LI line type that were either followed by at least one individual (colored) or not followed by any other bees (black). (D) Proportion of dances by LI line type that were followed (from panel B) broken down by LI of the follower. (E) Relationship between number of followers and duration of a dance by line. Point and line colors indicate LI of dancer. Best fit line represents the GLM, shaded area represents the 95% confidence interval. (F) Rate of turns per second in a dance by line. The large black dot in the box is the mean, the box is 25-75% of the data, whiskers represent 95% of the data. The violin shapes illustrate distribution of the data. Different letters above violins indicate statistically significant differences according to a post hoc Tukey test.

Although the high LI individuals danced less often, high LI dances had significantly more followers compared to low and control bees (Chi-square test: χ^2^= 13.93, df=2, p<0.001; Figure 4C). Low LI bees performed more dances that had no followers compared to high LI and control dances. We did not detect a statistically significant difference between the proportion of individuals from each line that followed each line of dancer (Chi-square test: χ^2^= 7.05, df = 4, p= 0.13, Figure 4D). Low LI individuals spent more time dancing; however they attracted fewer followers than high and control dancers, indicated by the significant interaction between the LI of the dancer and dance duration when predicting the number of followers (GLMM: χ^2^= 6.42, df=2, p=0.04; Figure 4E).

The relative attraction of dances of high LI bees could be due to the intensity of the dance. High LI bees performed more turns per second during their dances (ANOVA: χ^2^=12.8, df=2, p=0.002; Figure 4F). High LI dancers performed an average of 0.59 turns per second, significantly higher than low LI dancers, who performed an average of 0.52 turns per second (Tukey: t=-3.13, p=0.005). Control bees also performed more turns per second than low LI bees (Tukey: t=-2.5, p=0.03), but not different than high LI bees, at an average 0.62 turns per second (Tukey: t=-0.7, p=0.7).

## DISCUSSION

By combining techniques from experimental psychology and behavioral ecology, we have developed a system for investigating how variation in individual learning behavior drives collective cognition. We utilized this system to demonstrate that a laboratory-selected heritable learning behavior with natural individual variation scales to shape the collective performance of a honey bee colony on foraging tasks. In the lab, high LI honey bees learn to ignore familiar odors that they experienced without reinforcement, while readily learning novel odors. When a stimulus is rewarding, high LI bees exhibit increased attention to that information. One interpretation of reduced learning to a familiar, unrewarding, stimulus is that pre-exposure reduces attention to, and thus associability of, that stimulus. This interpretation is an extension of conditioned attention theory(29, 30), which proposes that latent inhibition is induced by allowing animals to focus their attention on important information(30–32). Our observations of field behavior of low and high LI individuals and colonies are consistent with this interpretation, whereby high LI individuals have stronger attention capacities to food compared to low LI individuals. Once high LI individuals have found a food location, they continue to revisit it, ‘attending’ more strongly to reinforced feeders over new ones. The increased impact of the resource on these bees could translate into stronger, more vigorous dances. In contrast, low LI individuals learn and visit both known and new feeders equally, dividing their attention across resources and acting more like generalist foragers. In mixed colonies, this broadened attention by low LI individuals may therefore make them the perfect audience for the high LI dancers, driving them to prefer feeders that high LI individuals preferentially visit. Under natural conditions, where queens mate with many different drones, most colonies would possess both types of learners, perhaps more closely resembling our mixed colonies(33). Attention is critical for many individual behaviors, including finding the correct mate or prey(34). We propose that this diversity of ‘attention’ aspect of individual cognitive phenotypes may enhance the overall efficacy with which a group finds and exploits resources(35). In summary, our work indicates that individual cognition scales to shape the collective cognition of animals solving critical ecologically relevant tasks.

## MATERIALS AND METHODS

### Obtaining queens and drones

To obtain queens for producing selected lines of a specific LI behavior, we performed the LI assay as outlined in(23, 24) on mature virgin queens 10 days after emergence. Briefly, we familiarized bees to an odor by puffing it at them 40 times every 5 minutes, then used the proboscis extension reflex to test their ability to learn to associate a food reward to the familiar versus a novel odor. Tested queens were placed into individually labelled queen cages and returned to a queenless colony until insemination, which typically occurred within a week of testing. To obtain fertile drones, we collected them from their returning unsuccessful mating flights at the entrance of colonies. We placed them into cages overnight in a queenless colony for LI testing the next day. After testing, drones were marked for individual identification and placed into a cage and placed into a queen bank for no longer than 3 days until inseminations occurred.

### Queen Inseminations

We used instrumental insemination to inseminate a queen with sperm from a single drone. We inseminated a high LI queen with a high LI drone, and a low LI queen with a low LI drone(36, 37). LI varies across individuals. However, for this behavioral selection, we used the highest and lowest LI scoring individuals to create the high and low colonies, respectively. We then introduced queens to small queenless colonies, then allowed the queens to produce workers for 1 month. Colonies were checked weekly to eliminate the possibility of supersedure.

### Cohoused Worker Preparation and Testing

To test the LI of foragers from each LI line, we placed frames of capped pupae from 3 high and 3 low LI colonies into 34°C incubators for 18 hours. After 18 hours, we used water based acrylic paint pens (Montana brand) to mark the abdomens of the eclosed bees with a color indicating their natal colony. Half of the bees were then returned to their natal colony and half were placed into an established control colony of an open-mated queen. Fewer bees were recovered from the established colony as many are recognized as non-nestmates and rejected. After 2 weeks, colonies were monitored every day until marked bees began to forage, ∼21 days after emergence. Returning nectar foragers were collected and tested for LI.

### Field Colony Experimental Setup

To explore the colony-level foraging behavior of the LI lines, we set up 4 treatment colonies for each of the colony types: a high LI colony, a low LI colony, a 50/50 mixed colony, and a control. We ran weekly field experiments for 6 weeks. For ease of identification, we always marked individuals from high LI colonies red, orange, and pink, and individuals from low LI colonies green, blue, yellow, and white. We continued to mark emerging bees from the same frames until we had 650 bees to form a colony, which took typically 2-3 days. To achieve relatively normal conditions for typical honey bee behavior, we supplemented workers from an unselected colony (control bees), who were not marked. For colony set up, see Supplementary Table 1. Bees were then placed into 4 different treatment colonies consisting of ∼1300 bees: high plus controls, low plus controls, 50/50 mixed high/low plus controls, and only control colony. Bees were provided a honeycomb and remained inside for 5 days before being placed for field experimentation. We then placed nucleus colonies into outdoor flight cages (3.05m × 12m) and replaced the honeycomb frame with an empty frame to induce foraging the night before the experiment. Water was provided as needed. We ran high, low, mixed, and control colonies concurrently in 4 different tents.

We used a familiar and novel feeder foraging assay to characterize colony level foraging behavior(38). We placed a feeder with 1M sucrose on Day 1, which remained in the same location all week and became the ‘familiar’ feeder (Figure 2). We then placed one novel feeder in different locations each day (Day 2 (X), Day 3 (Y), and Day 4 (Z)). Feeders had unique colors and unique odors and remained consistent throughout the experiment (Supplementary Table 2).

### Mixed Colony Round Dance Preparation and Data Collection

To evaluate round dance behavior of each of the selected lines, we created 6 50/50 mixed colonies as detailed above. To induce foraging behavior, we placed the colonies in a climate controlled indoor room for 10 days to allow bees to age which increases foraging behavior. After 10 days, we then placed all bees from each colony into a two-frame observation hive with glass walls. All comb surfaces were visible. We video recorded round dance behavior using a Panasonic HC-WXF991K, starting the recording 15 minutes before feeders were placed in the flight cage. For distances from the colony entrance, see figure 4A. We followed the same feeder placement pattern across 4 days, from Monday to Thursday, in Figure 4A. Round dance data was then extracted visually from watching videos. We recorded the LI line of the dancer according to the color marking on her thorax color, the feeder she visited according to the color mark on her abdomen (which also distinguished her as having visited a feeder), duration of the round dance, the LI line of the round dance followers, and the number of turns in a dance during the first 20 s of the dance, or less if the dance ended before 20 s elapsed. As the feeders were less than 12 m away from the colony, bees performed round dances which lack distinct ‘runs’ and often have incomplete turns. Video watchers were blind to the thorax and abdomen color associations between LI line and feeder visitation, respectively.

### Data Analysis Methods

To test whether bees exhibited a similar LI score as their parents regardless of where they were housed after emergence, we used a generalized linear model. We used LI score as the response variable, which fit a log-linear distribution, so we used a gaussian family with a log link. Our fixed predictor variables were the line from which the bees originated (high or low) and the colony type that they were placed in after emergence (either their natal colony or a control colony).

To evaluate the effect of colony composition on colony-level foraging behavior to novel and familiar feeders, we performed a general linear model with a gaussian error distribution on number of visits, with line and feeder as fixed predictor variables, as well as the interaction between line and feeder. We performed a generalized linear model with a binomial error distribution with a logit link function on percent revisitation, as it was a proportion comparing the number of revisits divided by the total number of visits. Line and feeder were fixed predictor variables, as well as the interaction between the line and feeder.

To explore whether the selected LI line of a forager bee influenced which feeder it visited while in the mixed colony, we used a general linear model with a gaussian error distribution on number of visits, with year, selected line and feeder as a fixed predictor variables, as well as the interactions between these three. We did find a significant three-way interaction between year, selected line, and feeder, which we present in Supplementary Table 6. Therefore, we treat years independently and performed two different GLMs with selected line and feeder as our fixed predictor variables, as the workers from the different years came from a new set of selected queens and drones, colonies were in nucleus Langstroth hive boxes in 2017 but were placed in observation colonies in 2018, as well as differences in weather.

To compare the round dance behavior among the selected lines, we examined the effect of dancer selected line on the duration of the round dance, intensity of dancing, number of turns by dancers, and number of followers of each dance using generalized linear models. To analyze whether the duration of the round dance differed across the learning lines, the duration of the round dance response variable fit a log-normal distribution, so we used a generalized linear mixed model with a gaussian family and a log link. The LI of the dancer was our fixed predictor variable. To evaluate which lines attracted more dancers, we used a chi-square test to compare the proportion of dances that attracted no followers across the lines. To evaluate whether there were differences in the number of turns the dancers from each line performed, we used a linear mixed model because the response variable - the number of turns per second, was normally distributed. Finally, we used a negative binomial mixed regression model using the package MASS(39) to understand how duration of a dance and the LI of the dancer interacted to predict the number of followers. 159 dances out of 908 total dances had no followers, requiring a zero-inflated model approach. We analyzed only bees that had paint marks on their abdomens, ensuring that they had visited a feeder.

We used an Analysis of Deviance Wald chi-square test using the function Anova in the MASS(39) package to further evaluate the overall effect of each fixed predictor variable and interaction. We used the lme4 package(40) to perform these tests unless otherwise noted. Post hoc tests were performed to determine the relationships between the different levels of fixed predictor variables and their interactions using the package emmeans(41). We use R(42) for analysis.

## Data Availability

Data will be available on FigShare and code will be available on Github upon publication. Data and code available upon request by reviewers.

### Ethical Compliance

Honey bees (Apis mellifera) were used in this study. Queens (reproductive females) and drones (males) were behaviorally selected using lab assays to create selected lines of colonies. Worker honey bees (non-reproductive females) were tested in the field. All colonies were kept with typical honey bee practices. There was no ethics committee involved in approving the animal husbandry protocol.

## Acknowledgement

We thank S. Ohrt, E Sezen, N Kulkarni, and A Phillips for help with data collection. We thank C. Kwapich and D. Charbonneau for comments on drafts of this manuscript. This grant was funded by NIH NIGMS R01GM113967 to BHS, JG, & NPW, and NIH NIGMS F32GM126728 to CNC.

## SUPPMEMENTARY MATERIALS

**Table S1:**
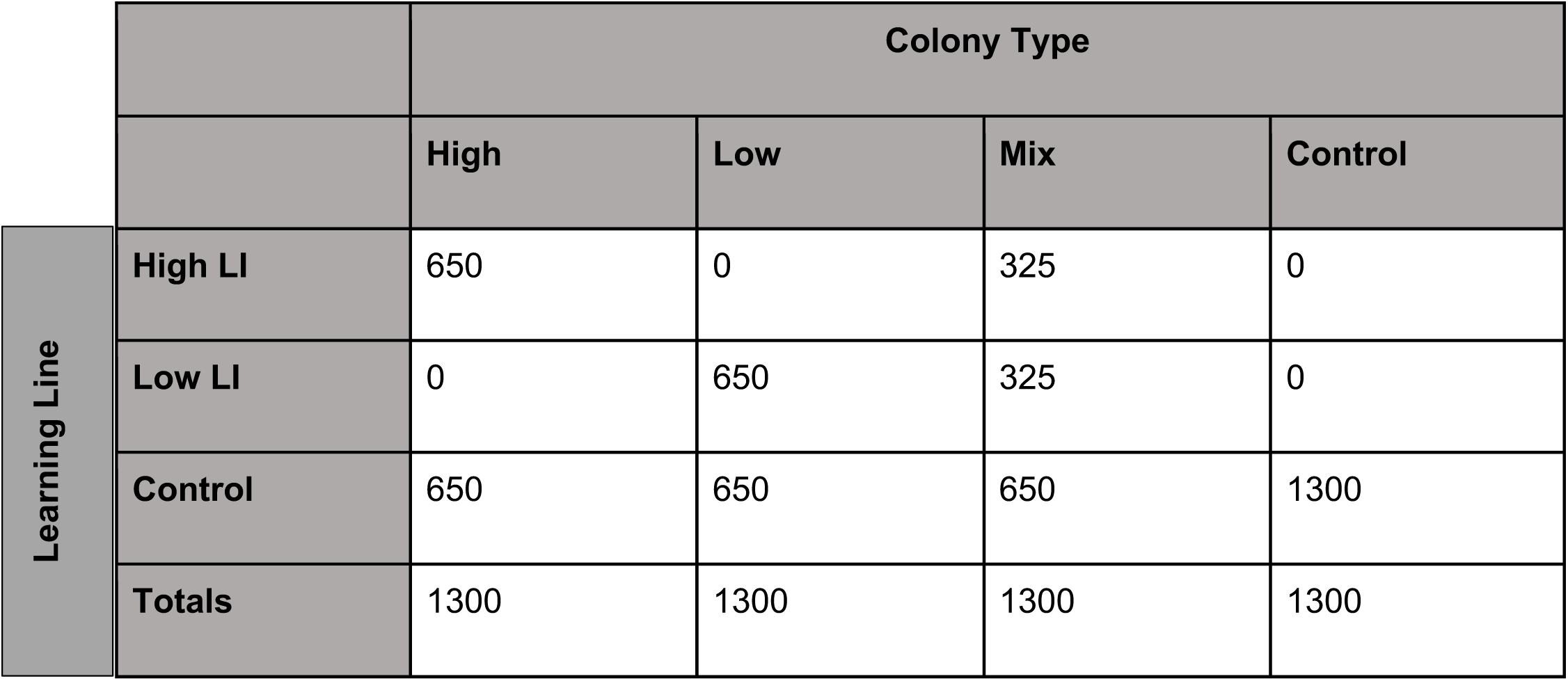
The number of honey bees in each experimental colony by genetic line. Each of the 4 created colonies were set up in this way each week. We counted and marked the thorax each bee from the learning lines, and counted but did not mark supplemental control bees.

**Table S2:**
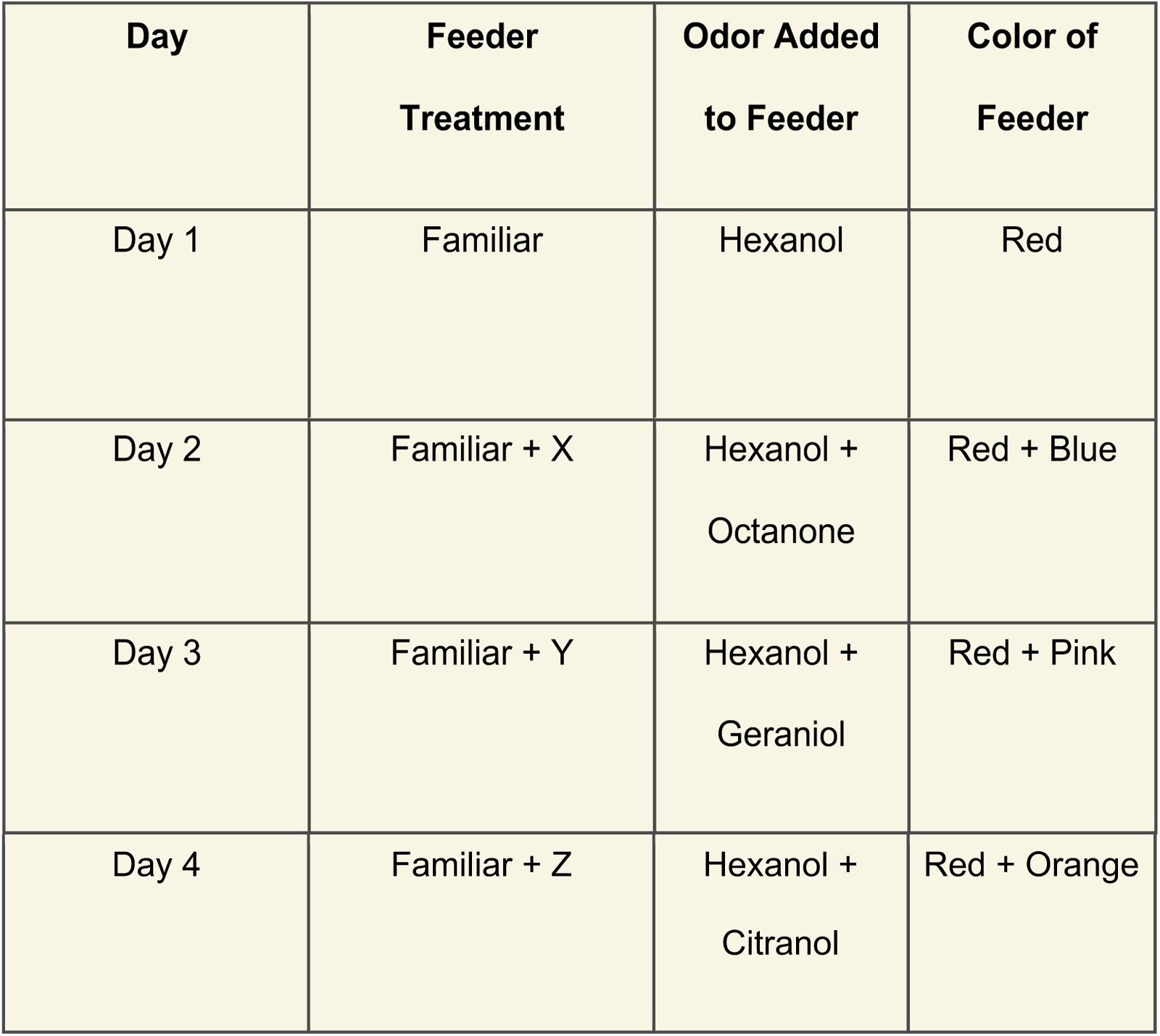
The weekly routine of feeder characteristics and placement. Each feeder had 1M sucrose solution. Color, odor, and location respectively varied by feeder. The treatment sequence was the same each week.

**Figure S1:**
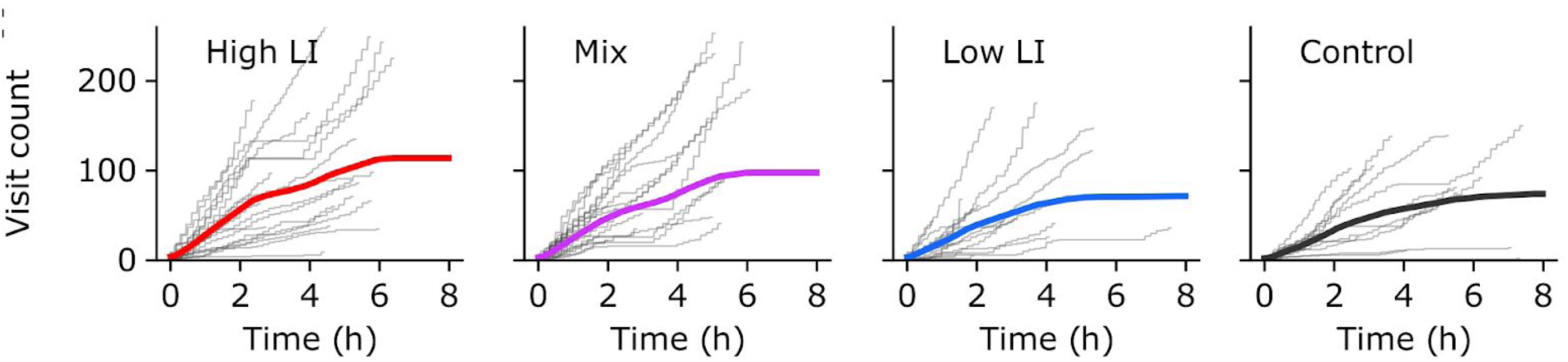
The cumulative visitation to all feeders over time, averaged across days. The thick colored line is the average, and the gray stepwise lines are visitation on a single day by a single colony. Colored lines are the same data shown in Figure 2B.

**Table S3:**
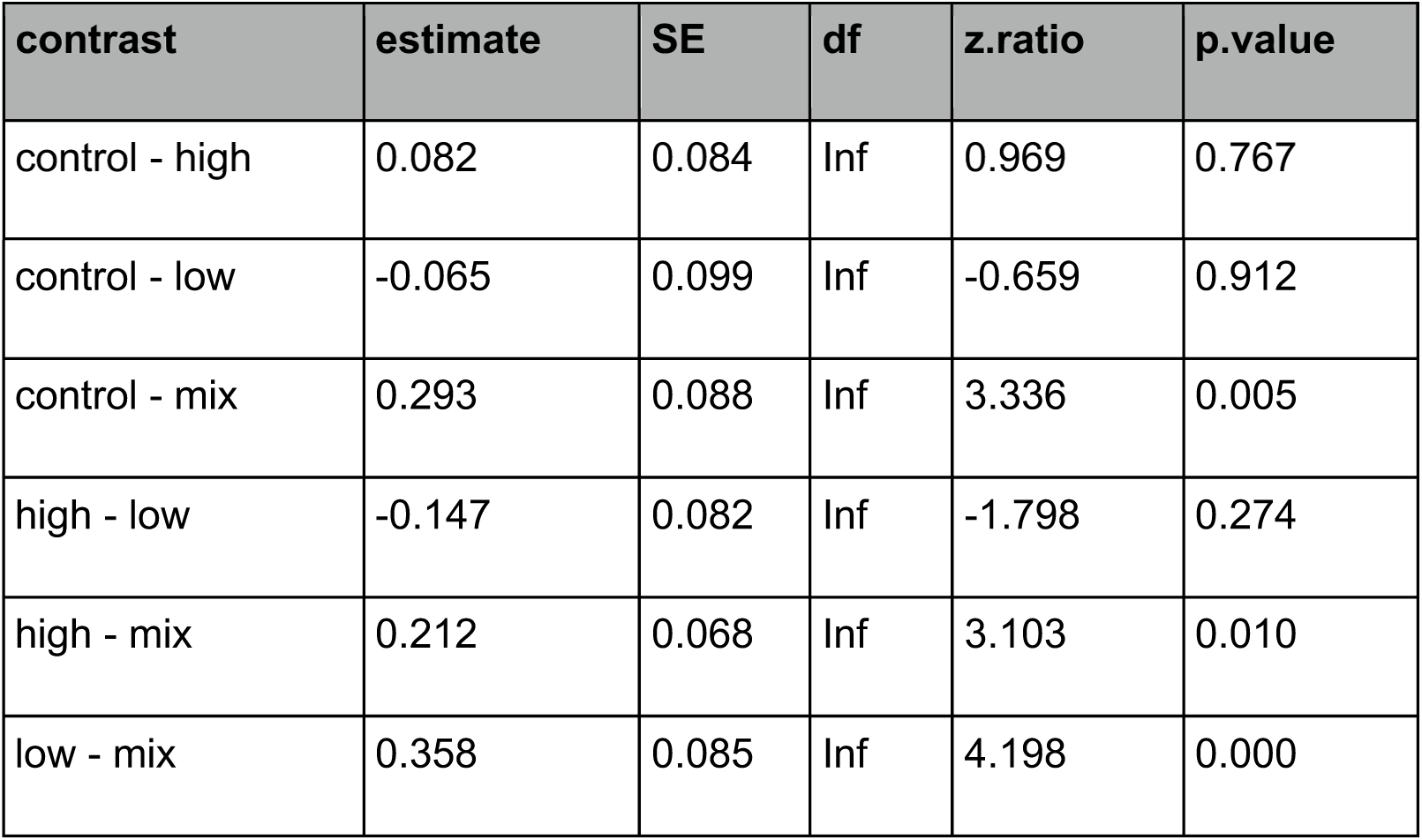
A table of the pairwise post hoc tests of how LI line predicts percent revisitation to all feeders, referenced in figure 2C.

**Table S4:**
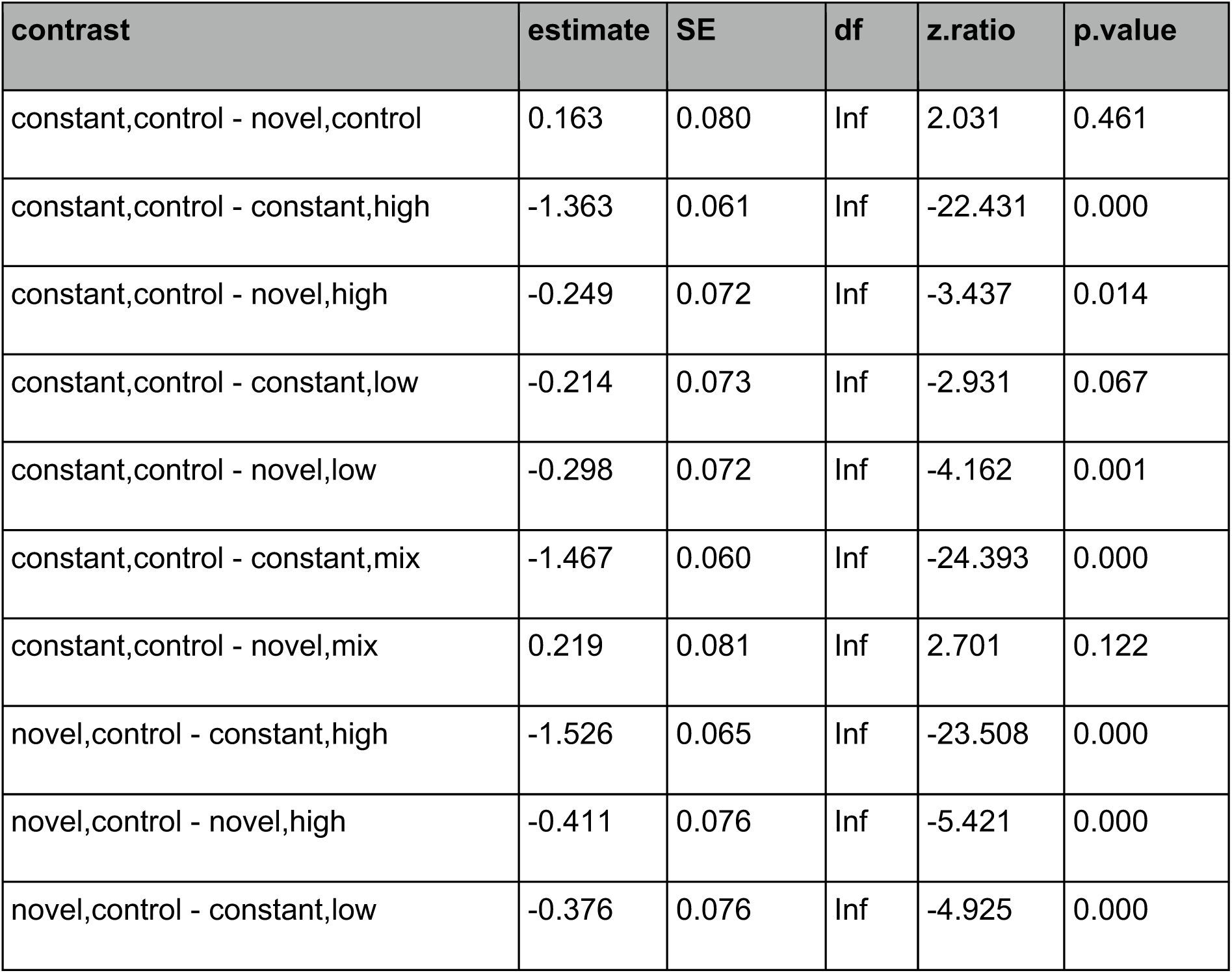
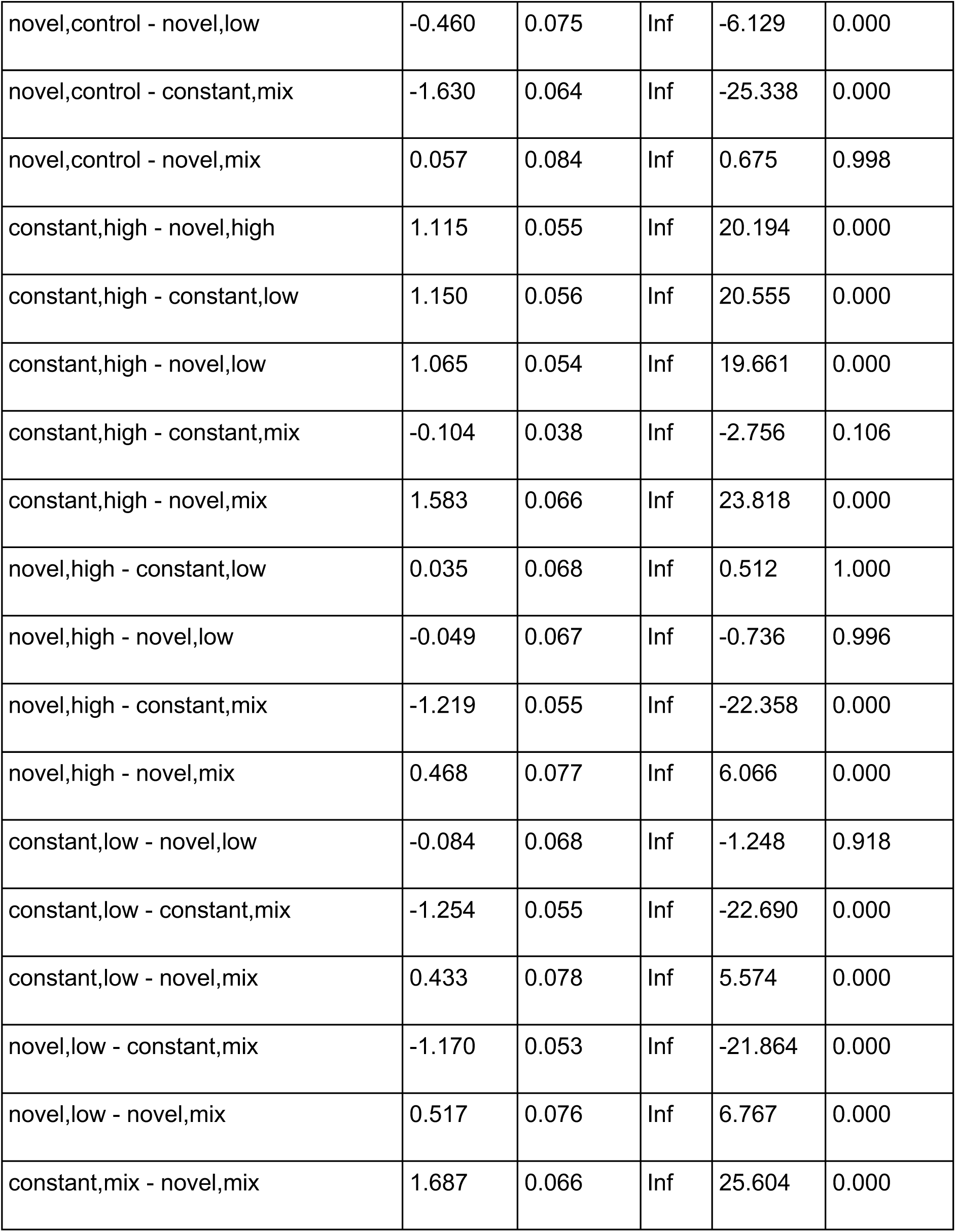
A table of the pairwise post hoc tests of how the Line*Feeder interaction predicts number of visits, which corresponds to letters in figure 2D.

**Table S5:**
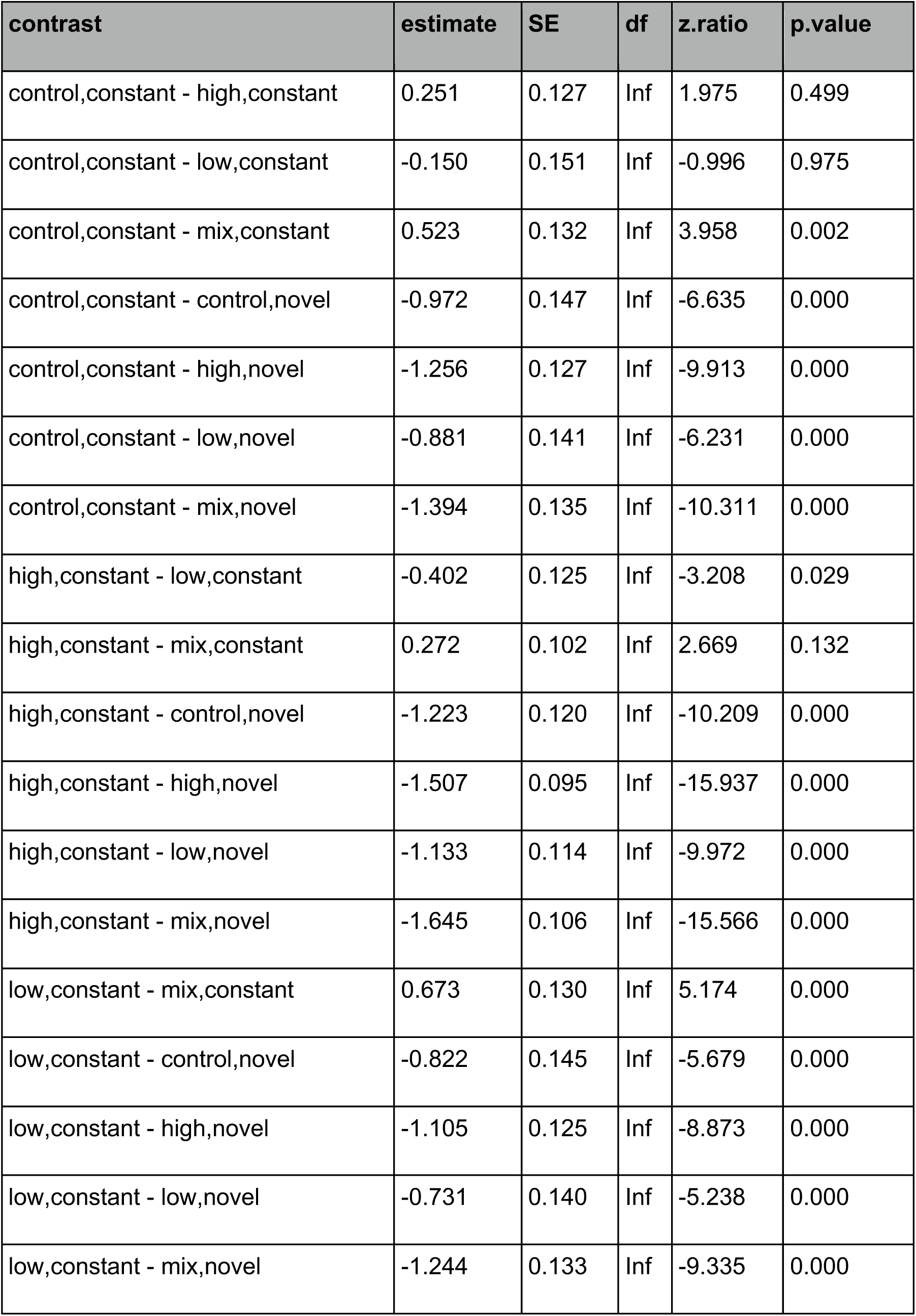
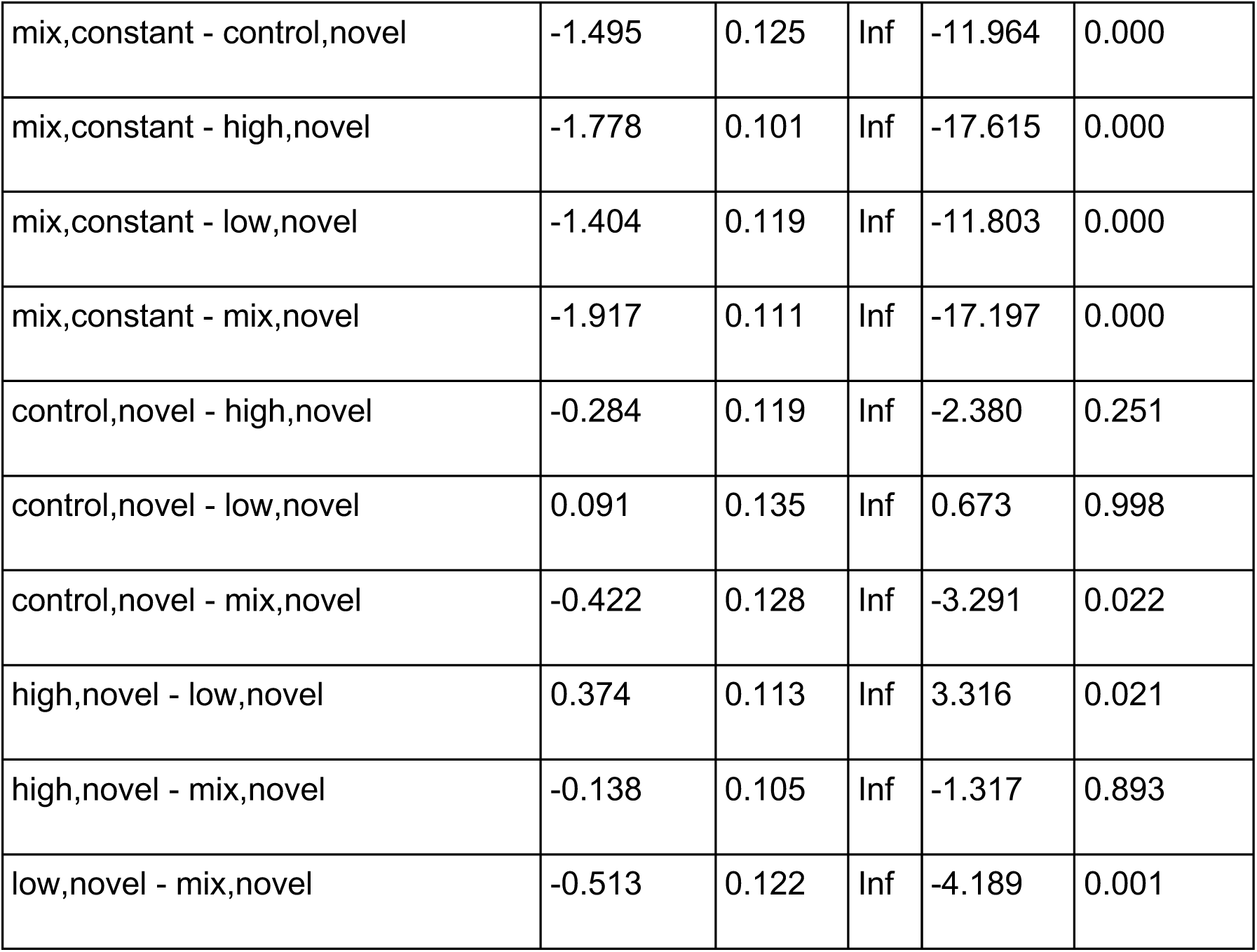
A table of the pairwise post hoc tests of how the Line*Feeder interaction predicts percent revisitation, which corresponds to letters in figure 2E.

**Table S6:**
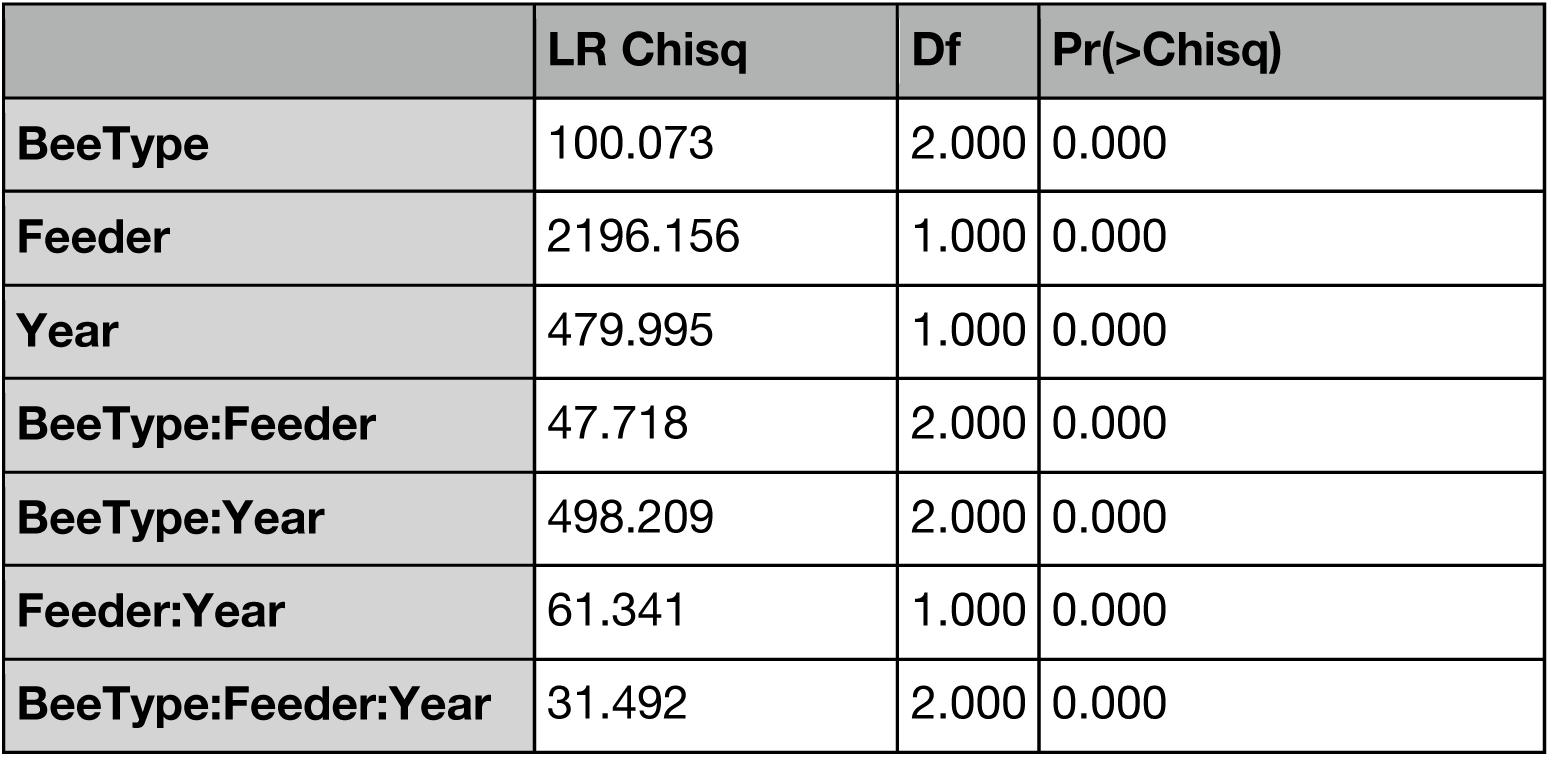
Individual visitation by bee type differed across two experimental years. GLM results showing the three-way interaction between year and the type of bee visiting a feeder (Figure 3). There is likely a difference in year because of several reasons, including 1) Colonies were selected from different queens from different breeders in 2017 and 2018 and climactic conditions were different in 2017 compared to 2018 even though experiments were done in the same time frame (June-July in 2017, June in 2018).

**Table S7:**
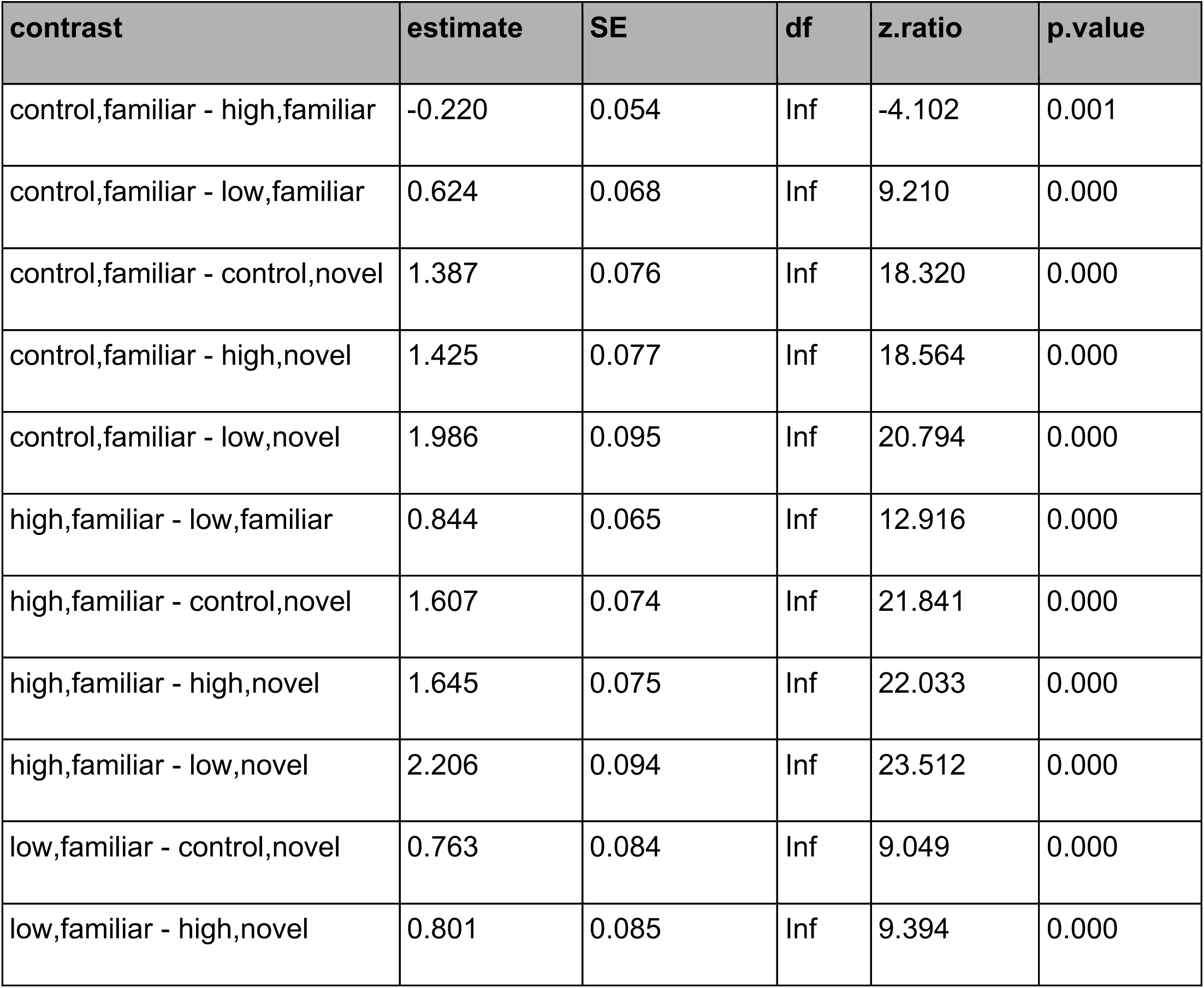
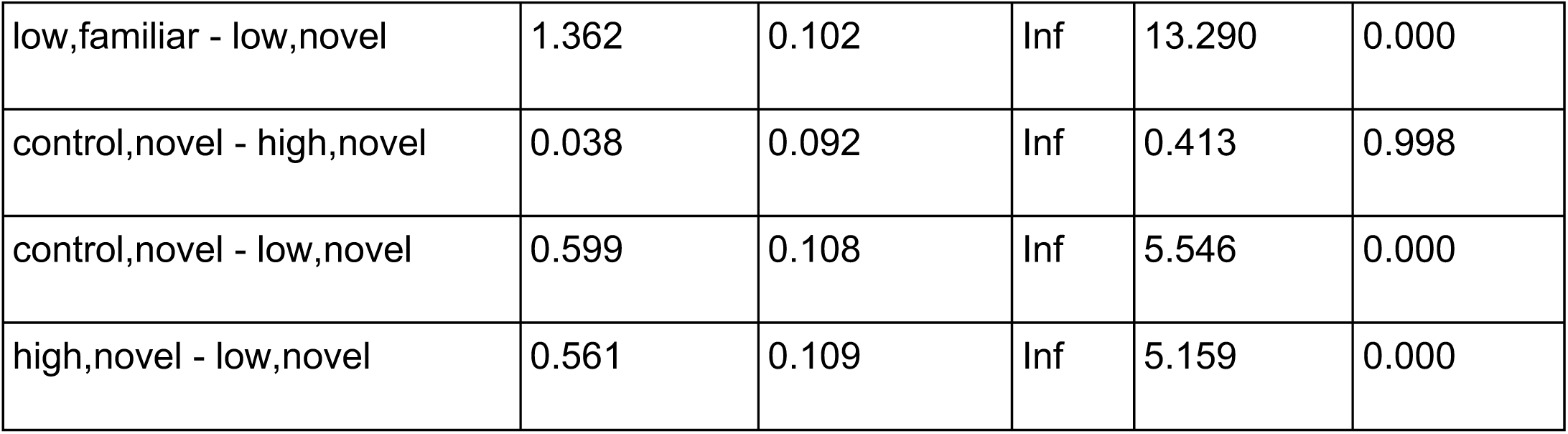
A table of the pairwise GLM contrasts of how the Line*Feeder interaction predicts number of visits by each line in the mixed colonies in 2017, which corresponds to letters in figure 3A.

**Table S8:**
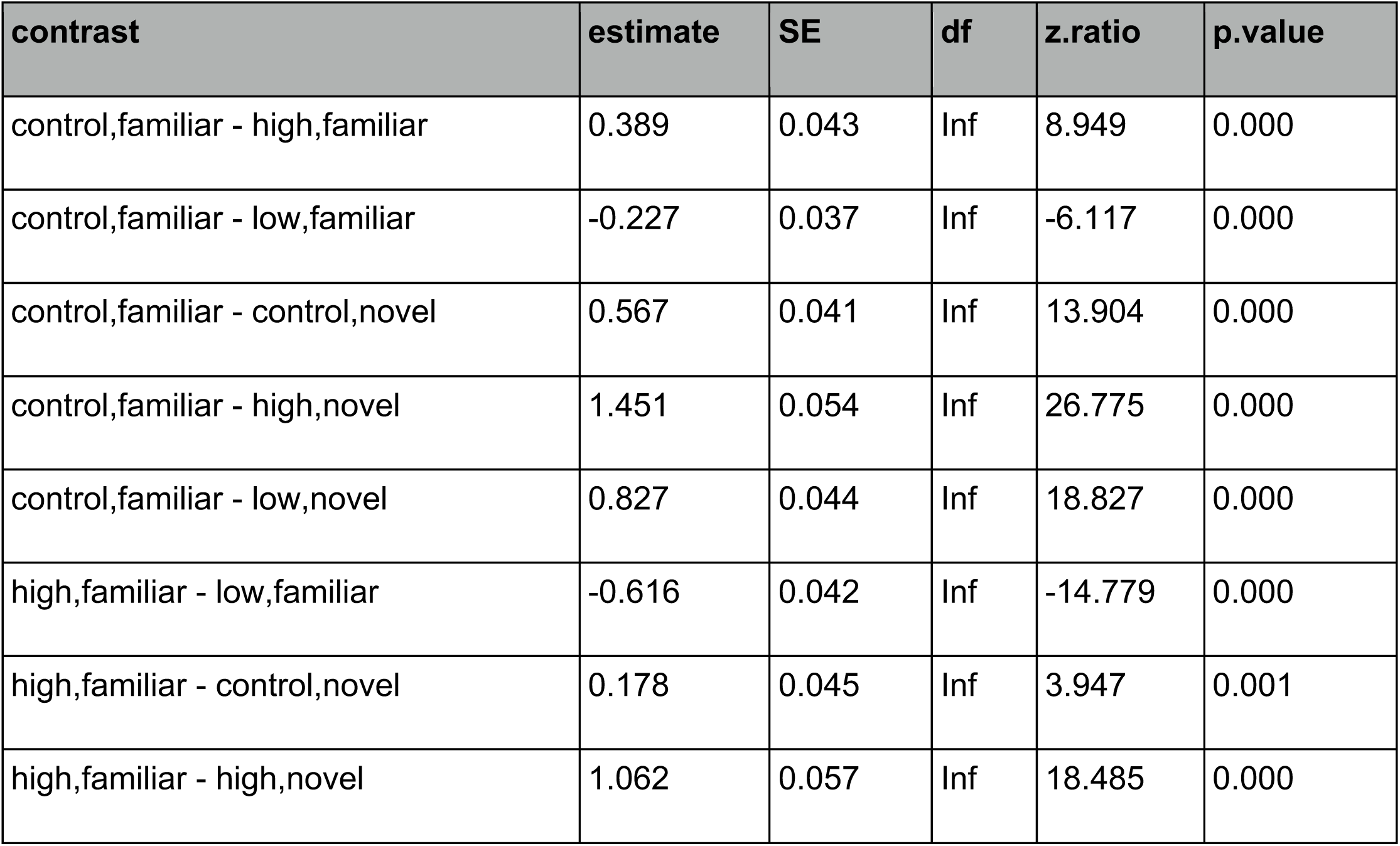
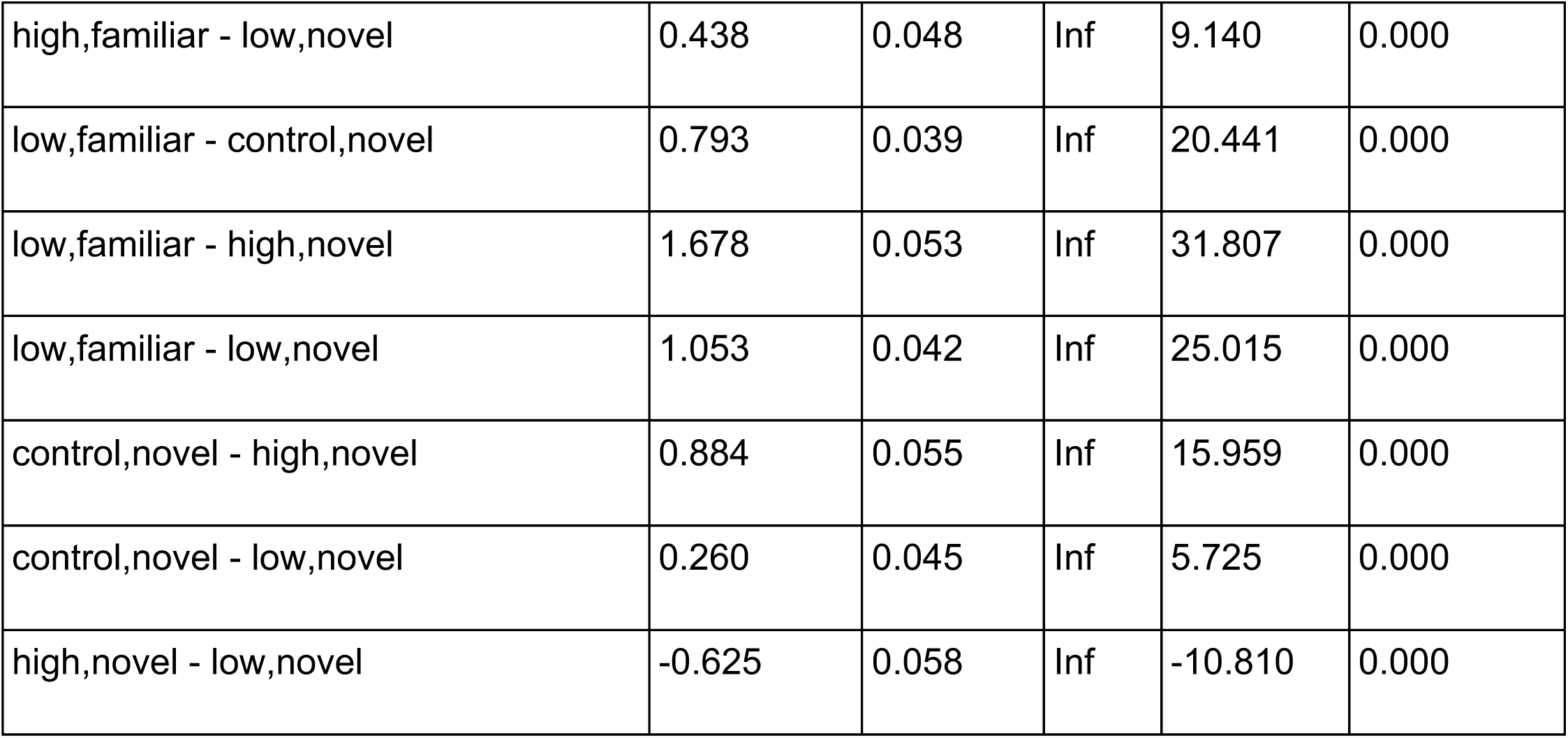
A table of the pairwise GLM contrasts of how the Line*Feeder interaction predicts number of visits by each line in the mixed colonies in 2018, which corresponds to letters in figure 3B.

