## Supplemental Materials for "Individual Learning Phenotypes Drive Collective Cognition"

**SUPPMEMENTARY MATERIALS - Individual Learning Phenotypes Drive Collective Cognition**

^†^Co-Senior Authors

**Learning Line**

|  | **Colony Type** | | | |
| --- | --- | --- | --- | --- |
|  | **High** | **Low** | **Mix** | **Control** |
| **High LI** | 650 | 0 | 325 | 0 |
| **Low LI** | 0 | 650 | 325 | 0 |
| **Control** | 650 | 650 | 650 | 1300 |
| **Totals** | 1300 | 1300 | 1300 | 1300 |

Table S1: **The number of honey bees in each experimental colony by genetic line.** Each of the 4 created colonies were set up in this way each week. We counted and marked the thorax each bee from the learning lines, and counted but did not mark supplemental control bees.

| **Day** | **Feeder**  **Treatment** | **Odor Added to Feeder** | **Color of Feeder** |
| --- | --- | --- | --- |
| Day 1 | Familiar | Hexanol | Red |
| Day 2 | Familiar + X | Hexanol + Octanone | Red + Blue |
| Day 3 | Familiar + Y | Hexanol + Geraniol | Red + Pink |
| Day 4 | Familiar + Z | Hexanol + Citranol | Red + Orange |

**Table S2: The weekly routine of feeder characteristics and placement.** Each feeder had 1M sucrose solution. Color, odor, and location respectively varied by feeder. The treatment sequence was the same each week.

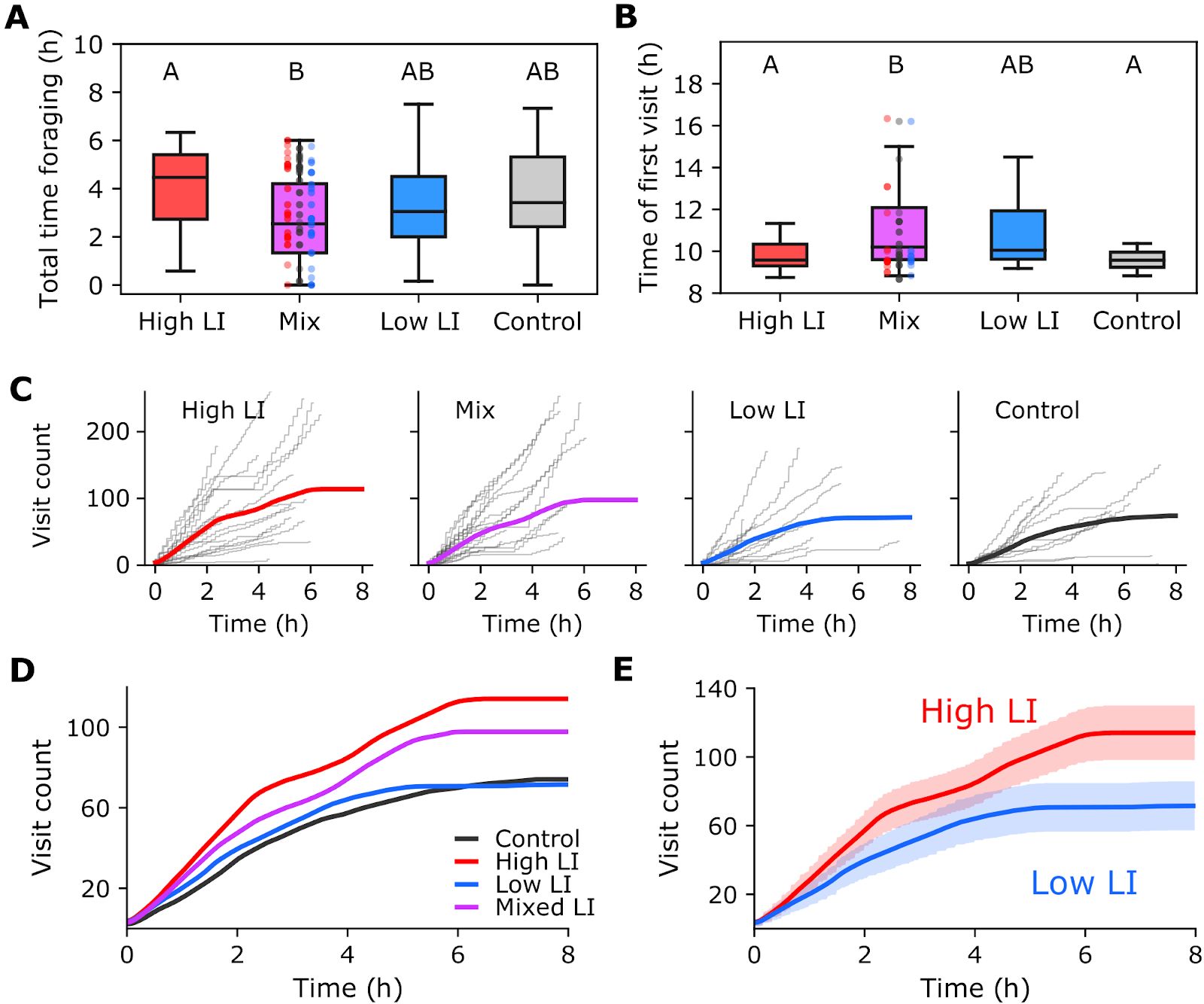

**Figure S1: The cumulative visitation to all feeders over time, averaged across days.** The thick colored line is the average, and the gray stepwise lines are visitation on a single day by a single colony. Colored lines are the same data shown in Figure 2B.

| **contrast** | **estimate** | **SE** | **df** | **z.ratio** | **p.value** |
| --- | --- | --- | --- | --- | --- |
| control - high | 0.082 | 0.084 | Inf | 0.969 | 0.767 |
| control - low | -0.065 | 0.099 | Inf | -0.659 | 0.912 |
| control - mix | 0.293 | 0.088 | Inf | 3.336 | 0.005 |
| high - low | -0.147 | 0.082 | Inf | -1.798 | 0.274 |
| high - mix | 0.212 | 0.068 | Inf | 3.103 | 0.010 |
| low - mix | 0.358 | 0.085 | Inf | 4.198 | 0.000 |

Table S3: A table of the pairwise post hoc tests of how LI line predicts percent revisitation to all feeders, referenced in figure 2C.

| **contrast** | **estimate** | **SE** | **df** | **z.ratio** | **p.value** |
| --- | --- | --- | --- | --- | --- |
| constant,control - novel,control | 0.163 | 0.080 | Inf | 2.031 | 0.461 |
| constant,control - constant,high | -1.363 | 0.061 | Inf | -22.431 | 0.000 |
| constant,control - novel,high | -0.249 | 0.072 | Inf | -3.437 | 0.014 |
| constant,control - constant,low | -0.214 | 0.073 | Inf | -2.931 | 0.067 |
| constant,control - novel,low | -0.298 | 0.072 | Inf | -4.162 | 0.001 |
| constant,control - constant,mix | -1.467 | 0.060 | Inf | -24.393 | 0.000 |
| constant,control - novel,mix | 0.219 | 0.081 | Inf | 2.701 | 0.122 |
| novel,control - constant,high | -1.526 | 0.065 | Inf | -23.508 | 0.000 |
| novel,control - novel,high | -0.411 | 0.076 | Inf | -5.421 | 0.000 |
| novel,control - constant,low | -0.376 | 0.076 | Inf | -4.925 | 0.000 |
| novel,control - novel,low | -0.460 | 0.075 | Inf | -6.129 | 0.000 |
| novel,control - constant,mix | -1.630 | 0.064 | Inf | -25.338 | 0.000 |
| novel,control - novel,mix | 0.057 | 0.084 | Inf | 0.675 | 0.998 |
| constant,high - novel,high | 1.115 | 0.055 | Inf | 20.194 | 0.000 |
| constant,high - constant,low | 1.150 | 0.056 | Inf | 20.555 | 0.000 |
| constant,high - novel,low | 1.065 | 0.054 | Inf | 19.661 | 0.000 |
| constant,high - constant,mix | -0.104 | 0.038 | Inf | -2.756 | 0.106 |
| constant,high - novel,mix | 1.583 | 0.066 | Inf | 23.818 | 0.000 |
| novel,high - constant,low | 0.035 | 0.068 | Inf | 0.512 | 1.000 |
| novel,high - novel,low | -0.049 | 0.067 | Inf | -0.736 | 0.996 |
| novel,high - constant,mix | -1.219 | 0.055 | Inf | -22.358 | 0.000 |
| novel,high - novel,mix | 0.468 | 0.077 | Inf | 6.066 | 0.000 |
| constant,low - novel,low | -0.084 | 0.068 | Inf | -1.248 | 0.918 |
| constant,low - constant,mix | -1.254 | 0.055 | Inf | -22.690 | 0.000 |
| constant,low - novel,mix | 0.433 | 0.078 | Inf | 5.574 | 0.000 |
| novel,low - constant,mix | -1.170 | 0.053 | Inf | -21.864 | 0.000 |
| novel,low - novel,mix | 0.517 | 0.076 | Inf | 6.767 | 0.000 |
| constant,mix - novel,mix | 1.687 | 0.066 | Inf | 25.604 | 0.000 |

Table S4: A table of the pairwise post hoc tests of how the Line*Feeder interaction predicts number of visits, which corresponds to letters in figure 2D.

| **contrast** | **estimate** | **SE** | **df** | **z.ratio** | **p.value** |
| --- | --- | --- | --- | --- | --- |
| control,constant - high,constant | 0.251 | 0.127 | Inf | 1.975 | 0.499 |
| control,constant - low,constant | -0.150 | 0.151 | Inf | -0.996 | 0.975 |
| control,constant - mix,constant | 0.523 | 0.132 | Inf | 3.958 | 0.002 |
| control,constant - control,novel | -0.972 | 0.147 | Inf | -6.635 | 0.000 |
| control,constant - high,novel | -1.256 | 0.127 | Inf | -9.913 | 0.000 |
| control,constant - low,novel | -0.881 | 0.141 | Inf | -6.231 | 0.000 |
| control,constant - mix,novel | -1.394 | 0.135 | Inf | -10.311 | 0.000 |
| high,constant - low,constant | -0.402 | 0.125 | Inf | -3.208 | 0.029 |
| high,constant - mix,constant | 0.272 | 0.102 | Inf | 2.669 | 0.132 |
| high,constant - control,novel | -1.223 | 0.120 | Inf | -10.209 | 0.000 |
| high,constant - high,novel | -1.507 | 0.095 | Inf | -15.937 | 0.000 |
| high,constant - low,novel | -1.133 | 0.114 | Inf | -9.972 | 0.000 |
| high,constant - mix,novel | -1.645 | 0.106 | Inf | -15.566 | 0.000 |
| low,constant - mix,constant | 0.673 | 0.130 | Inf | 5.174 | 0.000 |
| low,constant - control,novel | -0.822 | 0.145 | Inf | -5.679 | 0.000 |
| low,constant - high,novel | -1.105 | 0.125 | Inf | -8.873 | 0.000 |
| low,constant - low,novel | -0.731 | 0.140 | Inf | -5.238 | 0.000 |
| low,constant - mix,novel | -1.244 | 0.133 | Inf | -9.335 | 0.000 |
| mix,constant - control,novel | -1.495 | 0.125 | Inf | -11.964 | 0.000 |
| mix,constant - high,novel | -1.778 | 0.101 | Inf | -17.615 | 0.000 |
| mix,constant - low,novel | -1.404 | 0.119 | Inf | -11.803 | 0.000 |
| mix,constant - mix,novel | -1.917 | 0.111 | Inf | -17.197 | 0.000 |
| control,novel - high,novel | -0.284 | 0.119 | Inf | -2.380 | 0.251 |
| control,novel - low,novel | 0.091 | 0.135 | Inf | 0.673 | 0.998 |
| control,novel - mix,novel | -0.422 | 0.128 | Inf | -3.291 | 0.022 |
| high,novel - low,novel | 0.374 | 0.113 | Inf | 3.316 | 0.021 |
| high,novel - mix,novel | -0.138 | 0.105 | Inf | -1.317 | 0.893 |
| low,novel - mix,novel | -0.513 | 0.122 | Inf | -4.189 | 0.001 |

Table S5: A table of the pairwise post hoc tests of how the Line*Feeder interaction predicts percent revisitation, which corresponds to letters in figure 2E.

|  | **LR Chisq** | **Df** | **Pr(>Chisq)** |
| --- | --- | --- | --- |
| **BeeType** | 100.073 | 2.000 | 0.000 |
| **Feeder** | 2196.156 | 1.000 | 0.000 |
| **Year** | 479.995 | 1.000 | 0.000 |
| **BeeType:Feeder** | 47.718 | 2.000 | 0.000 |
| **BeeType:Year** | 498.209 | 2.000 | 0.000 |
| **Feeder:Year** | 61.341 | 1.000 | 0.000 |
| **BeeType:Feeder:Year** | 31.492 | 2.000 | 0.000 |

Table S6: **Individual visitation by bee type differed across two experimental years.** GLM results showing the three-way interaction between year and the type of bee visiting a feeder (Figure 3). There is likely a difference in year because of several reasons, including 1) Colonies were selected from different queens from different breeders in 2017 and 2018 and climactic conditions were different in 2017 compared to 2018 even though experiments were done in the same time frame (June-July in 2017, June in 2018).

| **contrast** | **estimate** | **SE** | **df** | **z.ratio** | **p.value** |
| --- | --- | --- | --- | --- | --- |
| control,familiar - high,familiar | -0.220 | 0.054 | Inf | -4.102 | 0.001 |
| control,familiar - low,familiar | 0.624 | 0.068 | Inf | 9.210 | 0.000 |
| control,familiar - control,novel | 1.387 | 0.076 | Inf | 18.320 | 0.000 |
| control,familiar - high,novel | 1.425 | 0.077 | Inf | 18.564 | 0.000 |
| control,familiar - low,novel | 1.986 | 0.095 | Inf | 20.794 | 0.000 |
| high,familiar - low,familiar | 0.844 | 0.065 | Inf | 12.916 | 0.000 |
| high,familiar - control,novel | 1.607 | 0.074 | Inf | 21.841 | 0.000 |
| high,familiar - high,novel | 1.645 | 0.075 | Inf | 22.033 | 0.000 |
| high,familiar - low,novel | 2.206 | 0.094 | Inf | 23.512 | 0.000 |
| low,familiar - control,novel | 0.763 | 0.084 | Inf | 9.049 | 0.000 |
| low,familiar - high,novel | 0.801 | 0.085 | Inf | 9.394 | 0.000 |
| low,familiar - low,novel | 1.362 | 0.102 | Inf | 13.290 | 0.000 |
| control,novel - high,novel | 0.038 | 0.092 | Inf | 0.413 | 0.998 |
| control,novel - low,novel | 0.599 | 0.108 | Inf | 5.546 | 0.000 |
| high,novel - low,novel | 0.561 | 0.109 | Inf | 5.159 | 0.000 |

Table S7: A table of the pairwise GLM contrasts of how the Line*Feeder interaction predicts number of visits by each line in the mixed colonies in 2017, which corresponds to letters in figure 3A.

| **contrast** | **estimate** | **SE** | **df** | **z.ratio** | **p.value** |
| --- | --- | --- | --- | --- | --- |
| control,familiar - high,familiar | 0.389 | 0.043 | Inf | 8.949 | 0.000 |
| control,familiar - low,familiar | -0.227 | 0.037 | Inf | -6.117 | 0.000 |
| control,familiar - control,novel | 0.567 | 0.041 | Inf | 13.904 | 0.000 |
| control,familiar - high,novel | 1.451 | 0.054 | Inf | 26.775 | 0.000 |
| control,familiar - low,novel | 0.827 | 0.044 | Inf | 18.827 | 0.000 |
| high,familiar - low,familiar | -0.616 | 0.042 | Inf | -14.779 | 0.000 |
| high,familiar - control,novel | 0.178 | 0.045 | Inf | 3.947 | 0.001 |
| high,familiar - high,novel | 1.062 | 0.057 | Inf | 18.485 | 0.000 |
| high,familiar - low,novel | 0.438 | 0.048 | Inf | 9.140 | 0.000 |
| low,familiar - control,novel | 0.793 | 0.039 | Inf | 20.441 | 0.000 |
| low,familiar - high,novel | 1.678 | 0.053 | Inf | 31.807 | 0.000 |
| low,familiar - low,novel | 1.053 | 0.042 | Inf | 25.015 | 0.000 |
| control,novel - high,novel | 0.884 | 0.055 | Inf | 15.959 | 0.000 |
| control,novel - low,novel | 0.260 | 0.045 | Inf | 5.725 | 0.000 |
| high,novel - low,novel | -0.625 | 0.058 | Inf | -10.810 | 0.000 |

Table S8: A table of the pairwise GLM contrasts of how the Line*Feeder interaction predicts number of visits by each line in the mixed colonies in 2018, which corresponds to letters in figure 3B.
